# Lipidomic analysis of postprandial mesenteric lymph from wild type and *Dennd5b* knockout mice

**DOI:** 10.64898/2026.11.11.765940

**Authors:** Khaga R. Neupane, Alexander Karakashian, Olivia Hage, Clairity Voy, Ryan Barney, Min Liu, Scott M. Gordon

**Author notes:** **Corresponding Author:** Scott M. Gordon Ph.D., Associate Professor, University of Kentucky, Saha Cardiovascular Research Center, 741 South Limestone, BBSRB Room B349, Lexington, KY 40536-0509.

## Abstract

Chylomicrons are intestine-derived lipoproteins that transport dietary lipids from the intestine to the systemic circulation. Although the composition of chylomicrons may influence their peripheral metabolism and biological functions, the lipid composition of newly secreted chylomicrons remains incompletely characterized. In this study, we combined a conscious mesenteric lymph fistula model with mass spectrometry-based lipidomics to characterize changes in the intestinal lymph lipidome following duodenal lipid administration in wild type and *Dennd5b*-deficient mice, a model of impaired intestinal chylomicron secretion. In wild type mice, lipid administration significantly increased lymphatic triglyceride output and altered the abundance of multiple lipid species, including glycerolipids, glycerophospholipids, and sphingolipids. Notably, ceramides and hexosylceramides exhibited disproportionate increases relative to their abundance in the administered lipid. In contrast, *Dennd5b*-deficient mice exhibited a near-complete blunting of the postprandial lymphatic lipidomic response, with markedly attenuated export across multiple lipid classes. Genotype-dependent differences were also evident in baseline lymph, indicating that *Dennd5b* influences lymphatic lipid composition under both basal and postprandial conditions. Together, these findings demonstrate that the physiological impact of *Dennd5b* deficiency on intestinal lipid trafficking extends well beyond triglycerides, underscoring an essential role for *Dennd5b*-dependent chylomicron secretion in the coordinated lymphatic export of dietary and endogenous lipids.

**Highlights:**

- Duodenal lipid administration significantly alters the mesenteric lymph lipidome in wild type mice.
- The mesenteric lymph lipidome generally shifts to reflect the lipid content of the ingested lipid, however, sphingolipids are disproportionately increased.
- *Dennd5b* deficiency broadly impairs postprandial lymphatic export of multiple lipid classes.

## Introduction

The intestinal epithelium is responsible for uptake of dietary lipids, intracellular re-esterification and packaging into chylomicrons, and the subsequent secretion of chylomicrons to the intestinal lymph system (1–4). Chylomicrons entering the intestinal lymphatics are transported through the mesenteric lymphatic system to the thoracic duct, where they enter the venous circulation and deliver dietary lipids to peripheral tissues (5). The protein and lipid composition of newly secreted chylomicrons may influence their subsequent metabolism, including susceptibility to lipoprotein lipase-mediated lipolysis, hepatic remnant clearance, and delivery of bioactive lipid species to peripheral tissues (6–8). Chylomicrons have also been implicated in the development of atherosclerotic cardiovascular disease due to whole-particle uptake by the vascular endothelium (9–11).

Because chylomicrons transport a complex mixture of bioactive lipids in addition to triglyceride, detailed characterization of the lipid composition of newly secreted chylomicrons could yield novel insight into how these particles are assembled in the intestine and subsequently metabolized. Analysis of chylomicrons isolated from postprandial plasma likely does not reflect the initial composition of these particles due to rapid lipolysis by lipoprotein lipase and clearance of remnant particles. The conscious lymph fistula mouse model has emerged as a powerful tool for isolating intestine-derived chylomicrons prior to their entry into the circulation (12, 13). Thus, analysis of intestinal lymph can allow examination of native properties of newly secreted chylomicrons and may yield information about their composition before they are acted upon by lipases. To our knowledge, detailed lipidomic profiling of fasting and postprandial mesenteric lymph has not been reported.

The DENN (Differentially Expressed in Normal and Neoplastic tissues) domain-containing family of proteins are guanine nucleotide exchange factors. These proteins are best known for facilitating the activation of Rab proteins to facilitate intracellular vesicular trafficking (14, 15). We have previously reported that mice lacking *Dennd5b* exhibit significantly impaired absorption of dietary triglyceride due to impaired secretion of chylomicrons from the intestinal epithelium (16). As a consequence, *Dennd5b^-/-^* mice exhibit reduced body mass and are resistant to diet-induced obesity and atherosclerosis (17, 18). These studies used biochemical and enzymatic assays to quantify triglyceride malabsorption and a limitation of our previous studies was that absorption of other lipid species was not examined.

In the present study, we sought to characterize the lipid composition of mesenteric lymph from wild type and *Dennd5b^-/-^*mice before and after lipid feeding. We aimed to determine if dietary lipid malabsorption in *Dennd5b^-/-^* mice is specific to triglyceride, as previously reported, or if absorption of other lipid species is also impacted by *Dennd5b* deficiency. To accomplish this, we utilized a conscious lymph fistula procedure to collect mesenteric lymph before and after administration of a mixed lipid bolus into the duodenum of wild type and *Dennd5b^-/-^* mice. Collected lymph samples were subjected to mass spectrometry-based lipidomic analysis, enabling detailed profiling of multiple lipid categories in baseline and post-bolus mesenteric lymph.

## Materials and Methods

### Mouse housing

Wild type and *Dennd5b^-/-^* mice were bred and maintained at the University of Kentucky Division of Laboratory Animals Resources in individually ventilated cages, with a maximum of 5 mice per cage. The mice were housed with normal day light: dark cycle, and the room temperature was maintained at 22°C (72°F). Teklad bedding system (#7090A, Harlan Teklad) was used in cages and mice were maintained on a standard chow diet (#2918 Envigo) with unlimited access to food and water. Adult mice were transferred to the University of Cincinnati Mouse Metabolic Phenotyping Center for the lymph fistula procedure. Institutional Animal Care and Use Committees at the University of Kentucky and the University of Cincinnati granted approval for all animal studies.

### Lymph collection

Mouse mesenteric lymph collection was performed at the University of Cincinnati, Mouse Metabolic Phenotyping Center. Mesenteric lymph was collected from *Dennd5b^+/+^* and *Dennd5b^-/-^*8-month-old male mice using the conscious lymph fistula model (19). Briefly, this procedure involves double cannulation surgery: the first cannula was inserted into the superior mesenteric lymph duct, and the second was placed in the duodenum 2 cm distal to the pylorus. Baseline lymph was collected for 1 hour prior to administration of the Intralipid™ (Sigma Aldrich, #I141) bolus. Mice then received the lipid emulsion (300 µL), composed primarily of soybean oil and egg phospholipids, followed by a continuous infusion of glucose and saline. Post-bolus lymph was collected continually for 5 hours. Collection tubes were changed hourly to permit analysis of 1-hour intervals. Lymphatic triglyceride content was measured using L-Type Triglyceride M colorimetric enzymatic assay (FujiFilm, #994-02891 and #990-02991).

### Lipidomic analysis of mesenteric lymph

#### Lymphatic lipid extraction

Baseline and post-bolus lymph collected between hours 2 and 3 after administration of the lipid bolus was used for lipidomic analysis. For each sample, 50 µL of lymph was mixed with 1 mL of extraction solvent (MTBE: MeOH = 3:1, v/v) containing an internal standard mixture and allowed to mix for 15 minutes. Next, 200 µL of ultrapure water was added and vortexed for 1 minute prior to centrifuging it at 12,000 rpm for 10 minutes. 200 µL of the upper organic layer was collected and evaporated using a vacuum concentrator. Reconstituted solution (acetonitrile: isopropanol=1:1, v/v) was used to dissolve the dry extract, and the resulting solution was subjected to ultra-performance liquid chromatography–tandem mass spectrometry (UPLC–MS/MS) platform consisting of a Nexera LC-40 UPLC system (Shimadzu, Japan) coupled to a QTRAP® 6500+ mass spectrometer (Sciex, USA).

#### Chromatography separation

Chromatographic separation was achieved using a Thermo Accucore™ C30 column (2.6 μm, 2.1 × 100 mm). The mobile phase consisted of solvent A (acetonitrile/water, 60:40, v/v) and solvent B (acetonitrile/isopropanol, 10:90, v/v), both supplemented with 0.1% formic acid and 10 mmol/L ammonium formate. The gradient elution program was as follows: 80:20 (v/v) at 0 min, 70:30 at 2 min, 40:60 at 4 min, 15:85 at 9 min, 10:90 at 14 min, 5:95 at 15.5 min, 5:95 at 17.3 min, 80:20 at 17.5 min, and 80:20 at 20 min. The flow rate was maintained at 0.35 mL/min, the column temperature was set to 45 °C, and the injection volume was 2 μL.

#### Mass spectrometry lipidomics

Mass spectrometric detection was performed using a QTRAP 6500+ triple quadrupole-linear ion trap mass spectrometer (QTRAP), QTRAP® 6500+ LC-MS/MS System, equipped with an electrospray ionization (ESI) Turbo Ion Spray interface, operating in positive and negative ion mode and controlled by Analyst v.1.6.3 software (Sciex). The ion source parameters were set as follows: source temperature 500 °C; ion spray voltage +5500 V (positive mode) and −4500 V (negative mode); curtain gas at 35 psi; ion source gas 1 (GS1) at 45 psi; and ion source gas 2 at 55 psi. Instrument tuning and mass calibration were achieved using polypropylene glycol solutions at concentrations of 10 μmol/L and 100 μmol/L for triple quadrupole and linear ion trap modes, respectively. Data acquisition was performed in multiple reaction monitoring (MRM) mode with nitrogen as the collision gas set to 5 psi. Declustering potential and collision energy parameters were optimized for individual MRM transitions. Specific MRM transitions were scheduled according to the retention time windows of lipid species eluting during defined chromatographic periods. Lipids were annotated based on their retention time and ion pair formation from MRM mode using the Innovation Omics inhouse database. Analyst v.1.6.3 (Sciex) was used for processing mass spectrum data.

### Statistical analyses

For comparisons between two groups, unpaired Student’s t-test was used. For comparisons of individual lipid class measurements by genotype and bolus status, two-way ANOVA with appropriate post-hoc testing for multiple comparisons. All statistical analyses were performed using GraphPad Prism and adjusted p values < 0.05 were regarded as statistically significant.

## Results

### *Dennd5b^-/-^* mice exhibit significantly reduced intestinal triglyceride output in response to a duodenal lipid bolus

To quantify intestinal lipid output, the conscious lymph fistula cannulation was performed on wild type (*Dennd5b^+/+^)* and *Dennd5b^-/-^* mice. Mesenteric lymph was collected at baseline and hourly after duodenal administration of an Intralipid bolus (**Figure 1A**). Triglyceride (TG) concentrations were measured by enzymatic assay in lymph collected at baseline and during the post-bolus period. A steady increase in cumulative intestinal triglyceride output was observed in wild type mice over the 5-hour post-bolus period (**Figure 1B**). *Dennd5b^-/-^* mice had significantly reduced intestinal triglyceride output compared to wild type mice (14.2 mg vs. 0.5 mg at 5 hours post-bolus, p = 0.007). This phenotype is consistent with our previous reports that *Dennd5b^-/-^* mice have a significantly blunted plasma triglyceride elevation after receiving gastric lipid gavage (16, 18).

**Figure 1.**
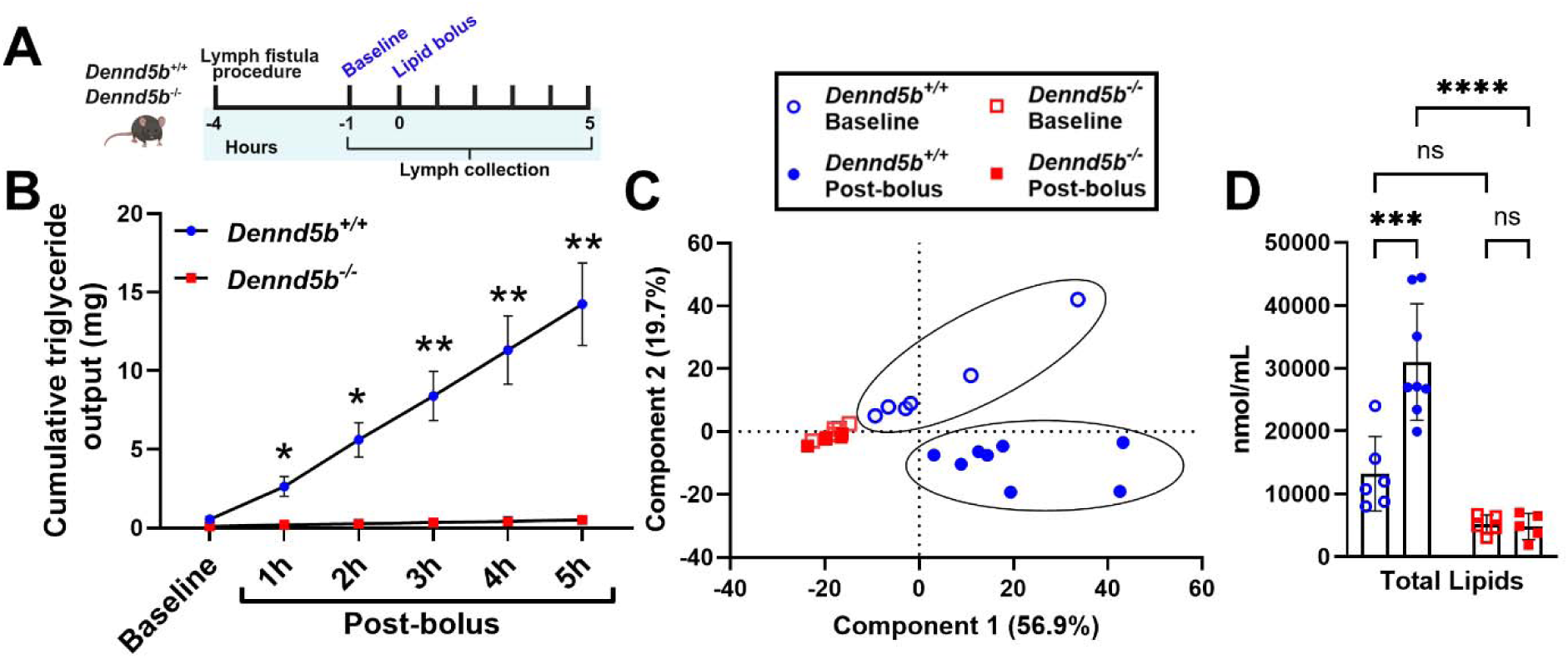
*Dennd5b* deficient mice exhibit significantly lower lymphatic lipid content after administration of a duodenal lipid bolus. (A) Schematic illustrating the experimental timeline for mouse mesenteric lymph collection. *Dennd5b^+/+^* and *Dennd5b^-/-^* mice underwent double cannulation lymph fistula surgery followed by collection of baseline and post-bolus lymph. (B) Cumulative lymphatic triglyceride (TG) output measured in *Dennd5b^+/+^* and *Dennd5b^-/-^* mice at baseline and during 5 hours post lipid bolus feeding. Data are mean +/- SEM (n=5-8/group). Comparison was made by two-way ANOVA with Šídák’s multiple comparisons test. (C) Principal component analysis (PCA) of lipidomic data obtained by analyzing mesenteric lymph collected at baseline and post-bolus conditions in *Dennd5b^+/+^* and *Dennd5b^-/-^* mice. (D) Lymphatic total lipid concentrations of the mesenteric lymph collected from *Dennd5b^+/+^* and *Dennd5b^-/-^* mice at baseline and 3 hours post-bolus. Data are presented as mean ± SEM. Comparisons were performed using two-way ANOVA with Tukey’s multiple-comparisons test. For all statistical analyses: *P < 0.05, **P < 0.01, ***P < 0.001, ****P < 0.0001, ns = not significant.

### *Dennd5b*-deficiency broadly affects intestinal lipid absorption in mice

To determine if genetic disruption of *Dennd5b* affects the absorption of other lipids, quantitative lipidomics was performed on baseline and post-bolus (collected during the 2 to 3 hour post-bolus interval) mesenteric lymph using an ultra-performance liquid chromatography tandem mass spectrometry (UPLC-MS/MS) detection platform. This analysis measured concentrations of 746 distinct lipid species. Principal component analysis indicated that there were differences in the lipid composition between *Dennd5b^+/+^* and *Dennd5b^-/-^* mice at baseline (**Figure 1C)**. Furthermore, this analysis revealed that the lymph lipidome in wild type mice experienced substantial compositional changes in response to the lipid bolus, however a similar lipidome shift did not occur in the post-bolus *Dennd5b^-/-^*lymph (**Figure 1C)**. Consistent with this, analysis of the total lipid content showed that *Dennd5b^+/+^* lymph had significantly increased total lipid content in response to the lipid bolus, however, the lipid content of *Dennd5b^-/-^*lymph did not change after the lipid bolus (**Figure 1D)**.

To examine the contributions of specific lipids to the observed effects on lymph lipid composition, quantified lipids were organized by lipid category and class. Table 1 presents the average concentrations by lipid class and fold-changes in response to the lipid bolus by genotype. In response to the lipid bolus, wild type lymph exhibited increased concentrations of several lipid categories including glycerolipids, sphingolipids, and some glycerophospholipids, whereas *Dennd5b^-/-^* lymph exhibited minimal or reduced responses for all lipids **(Table 1)**.

**Table 1.** Lipid concentrations in mesenteric lymph from *Dennd5b^+/+^* and *Dennd5b^-/-^* mice at baseline and after a lipid bolus. Values in “Baseline” and “Post-bolus” columns are lipid concentrations in nmol/mL. The “Fold-change” columns represent the fold change between post-bolus and baseline for each genotype. Color shading indicates the magnitude and direction of fold-change in response to lipid administration, with red denoting increased concentration and blue denoting decreased concentration relative to baseline for each genotype.

|  |  | *Dennd5b*<sup>+/+</sup> |  |  | *Dennd5b*<sup>-/-</sup> |  |  |
|---|---|---|---|---|---|---|---|
| Lipid category | Lipid class | Baseline | Post-bolus | Fold-change | Baseline | Post-bolus | Fold-change |
| GL (Glycerolipids) | TG (Triglyceride) | 6,947.9 | 16,847.9 | 2.4 | 2,389.7 | 2,402.7 | 1.0 |
|  | DG (Diglyceride) | 600.1 | 3,461.7 | 5.8 | 680.8 | 510.3 | 0.7 |
|  | MG (Monoglyceride) | 19.6 | 58.3 | 3.0 | 27.5 | 24.6 | 0.9 |
| GP (Glycerophospholipids) | LNAPE (Lys-N-acylphosphatidylethanolamine) | 13.1 | 72.9 | 5.6 | 2.1 | 3.7 | 1.8 |
|  | LPE (Lysophosphatidylethanolamine) | 88.5 | 85.4 | 1.0 | 72.8 | 49.5 | 0.7 |
|  | PE (Phosphatidylethanolamine) | 402.7 | 649.4 | 1.6 | 111.6 | 108.3 | 1.0 |
|  | LPC (Lysophosphatidylcholine) | 83.9 | 79.7 | 0.9 | 76.6 | 52.7 | 0.7 |
|  | PC (Phosphatidylcholine) | 1,845.1 | 2,714.7 | 1.5 | 593.5 | 580.2 | 1.0 |
|  | LPG (Lysophosphatidylglycerol) | 0.3 | 0.3 | 1.0 | 0.1 | 0.1 | 1.0 |
|  | PG (Phosphatidylglycerol) | 89.1 | 123.9 | 1.4 | 44.7 | 44.8 | 1.0 |
|  | LPI (Lysophosphatidylinositol) | 0.7 | 0.2 | 0.3 | 0.5 | 0.3 | 0.6 |
|  | PI (Phosphatidylinositol) | 213.6 | 252.6 | 1.2 | 64.9 | 68.7 | 1.1 |
|  | PS (Phosphatidylserine) | 94.7 | 299.7 | 3.2 | 15.3 | 25.8 | 1.7 |
|  | SM (Sphingomyelin ) | 11.0 | 8.2 | 0.7 | 5.0 | 4.2 | 0.8 |
|  | PA (Phosphatidic Acid) | 554.0 | 2,375.4 | 4.3 | 50.4 | 121.7 | 2.4 |
| FA (Fatty Acyls) | CAR (Acylcarnitines) | 1.6 | 1.7 | 1.1 | 1.0 | 0.7 | 0.7 |
|  | FFA (Free Fatty Acids) | 276.3 | 341.0 | 1.2 | 277.6 | 199.8 | 0.7 |
| SP (Sphingolipids) | Cer (Ceramide) | 1,129.8 | 3,069.7 | 2.7 | 378.8 | 339.8 | 0.9 |
|  | HexCer (Hexosylceramide) | 42.8 | 101.0 | 2.4 | 30.1 | 23.9 | 0.8 |
| ST (Sterol Lipids) | BA (Bile Acids) | 24.4 | 24.6 | 1.0 | 15.8 | 17.4 | 1.1 |
|  | CE (Cholesteryl Ester) | 262.2 | 66.6 | 0.3 | 113.2 | 74.7 | 0.7 |
|  | Cholesterol | 483.3 | 327.2 | 0.7 | 179.0 | 151.7 | 0.8 |
| PR (Prenol Lipids) | CoQ (Coenzyme Q) | 1.0 | 1.7 | 1.7 | 0.4 | 0.3 | 0.8 |
| **Total Lipid** |  | **13185.7** | **30964** | **2.3** | **5131.3** | **4805.9** | **0.9** |

### Lymphatic lipid composition analysis by relative molar concentrations

Molar percent composition analysis of mesenteric lymph revealed glycerolipids (GL) as the predominant lipid class across all conditions, with roughly similar molar distributions (∼60%) between genotypes **(Supplementary Table 1)**. In *Dennd5b^+/+^*mice, lipid administration shifted this distribution toward higher GL (from 57% at baseline to 66% post-bolus) at the expense of other classes, particularly glycerophospholipids (GP) (from 26% at baseline to 22% post-bolus) and sterol lipids (ST) (from 6 % at baseline to 1% post-bolus) **(Supplementary Table 1)**. Notably, the drop in mol % of sterols post-bolus is likely due to the relative absence of cholesteryl esters in the administered lipid solution. Other lipid classes including fatty acyls (FA), sphingolipids (SP), and prenol lipids (PR) remained low and showed only modest changes. In contrast, percent mol composition comparisons of lymph samples from *Dennd5b^-/-^*mice revealed little to no response to lipid feeding. Mol % composition of the Intralipid solution used for lipid bolus was: 73% GL, 21% GP, 0.6% FA, 4% SP, 0.002% ST, 0% PR, and 1.3 % others **(Supplementary Table 1)**.

### Post-bolus lymph generally resembles the molar composition of ingested lipid

To determine if the baseline to post-bolus change in lymph lipid composition reflects the lipid composition of the administered lipid bolus, we analyzed the mol % shifts of each lipid category in response to the lipid bolus in both *Dennd5b^+/+^* and *Dennd5b^-/-^*lymph. We graphed the baseline difference between Intralipid and baseline lymph (delta mol % for Intralipid vs baseline; x-axis) and the bolus effect (delta mol % for post-bolus vs baseline; y-axis). Thus, if the Intralipid composition is consistent with the post-bolus changes, the points should fall along a relationship with a slope ≈ 1 (**Figure 2**, green dotted lines). This analysis revealed that, for *Dennd5b^+/+^*mice, most lipid categories were distributed near the green diagonal and 5 of the 6 clearly changing lipid categories moved in the expected direction based on the Intralipid input (**Figure 2A**). Sphingolipids (SP) are the exception as they move in the opposite of the expected direction. For *Dennd5b^-/-^* mice, which have broadly impaired absorption of dietary lipids, data points for all lipid categories remained very close to the x-axis indicating that the lipid composition of the lymph did not change to reflect the composition of the administered lipid (**Figure 2B**). These results support the conclusion that a minimal level of each administered lipid type enters the lymph in *Dennd5b^-/-^* mice.

**Figure 2.**
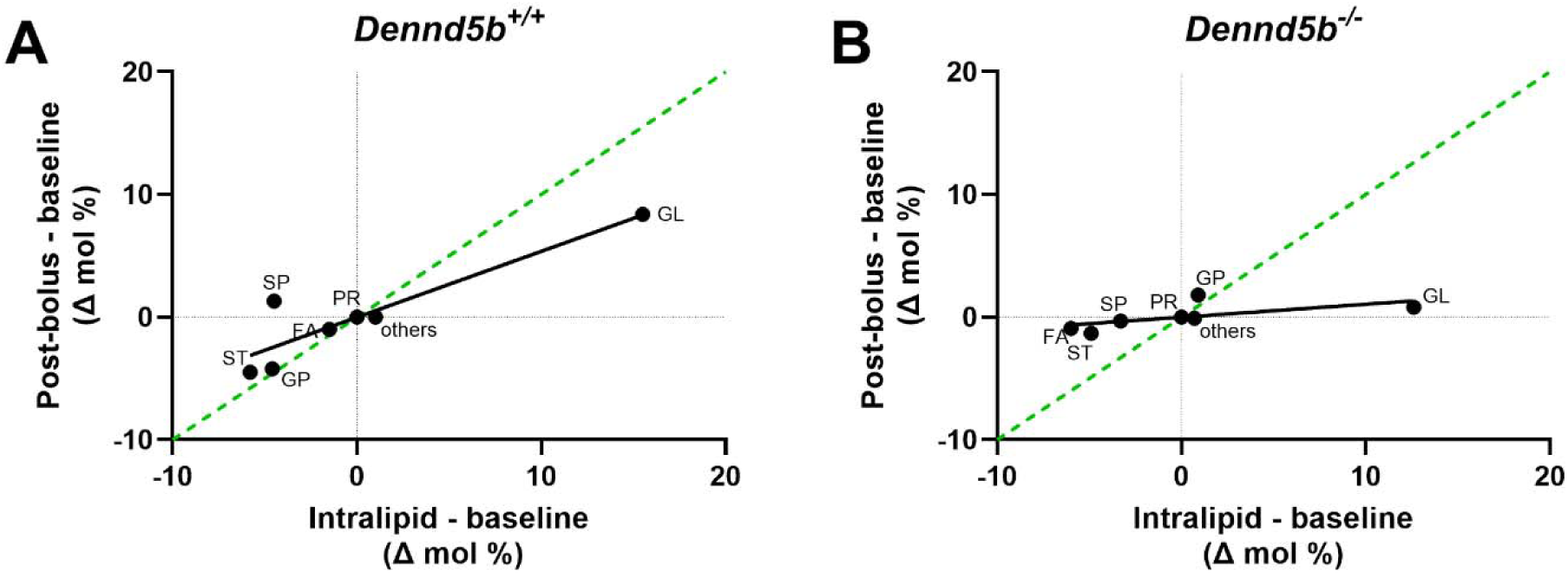
Post-bolus lymph lipidome shifts to resemble the molar composition of ingested lipid in *Dennd5b^+/+^* but not *Dennd5b^-/-^*mice. To determine if the shifts in lipid composition in response to the lipid bolus were consistent with the composition of the administered Intralipid, we calculated the difference in lipid class mole % between Intralipid and baseline lymph (x-axis, Intralipid difference) and the difference in lipid class mole % between post-bolus and baseline lymph (y-axis, bolus effect). (A-B) Scatter plots showing the relationship between post-bolus lipid class shifts and the composition of the administered lipid. The dotted green line represents a predicted direct relationship (slope = 1). Therefore, points close to the green line represent lipid classes whose post-bolus mole % shift was consistent with that lipid’s relative abundance in the administered lipid. This analysis was performed in (A) *Dennd5b^+/+^* and (B) *Dennd5b^-/-^* lymph. Each point represents a lipid class, including glycerolipids (GL), glycerophospholipids (GP), fatty acyls (FA), sphingolipids (SP), sterols (ST), prenol lipids (PR), and others.

### Analysis of specific lipid species concentrations in post-bolus lymph by lipid category

#### Glycerolipids

To determine the impact of lipid administration on lymphatic glycerolipid in *Dennd5b^+/+^* and *Dennd5b^-/-^*mice, total and individual glycerolipid classes were analyzed at baseline and following a lipid bolus **(Figure 3A–D)**. At baseline, wild type mice exhibited higher levels of total glycerolipid compared to *Dennd5b^-/-^*mice although this was not statistically significant **(Figure 3A)**. Following lipid bolus, total glycerolipid increased significantly in the lymph collected from *Dennd5b^+/+^* mice (2.7 fold; p < 0.0001), whereas no change was observed in *Dennd5b^-/-^* lymph. A similar pattern was observed for triglyceride (TG) and diglyceride (DG) **(Figure 3B, C)**, where *Dennd5b^+/+^* mice experienced significant increases after the bolus, while *Dennd5b^-/-^* mice maintained relatively low levels with no post-bolus increase. Monoglyceride (MG) levels **(Figure 3D)** were not different between wild type and *Dennd5b^-/-^* baseline lymph, but exhibited a moderate but significant increase following lipid bolus administration in wild type but not *Dennd5b^-/-^* mice. Differential analysis of individual glycerolipid species revealed that in the mesenteric lymph collected from *Dennd5b^+/+^* mice **(Figure 3E)**, concentrations of a large number of individual glycerolipids species were significantly increased following the bolus (116 species), with only a small subset whose concentrations decreased post-bolus (21 species). These effects were dominated by TG species with acyl chain lengths between 16 and 22 carbons, with some contributions from DG and MG species. In contrast, *Dennd5b^-/-^* mice **(Figure 3F)** exhibited a dramatically attenuated glycerolipid response, with only one species enriched post-bolus and three species decreasing post-bolus.

**Figure 3.**
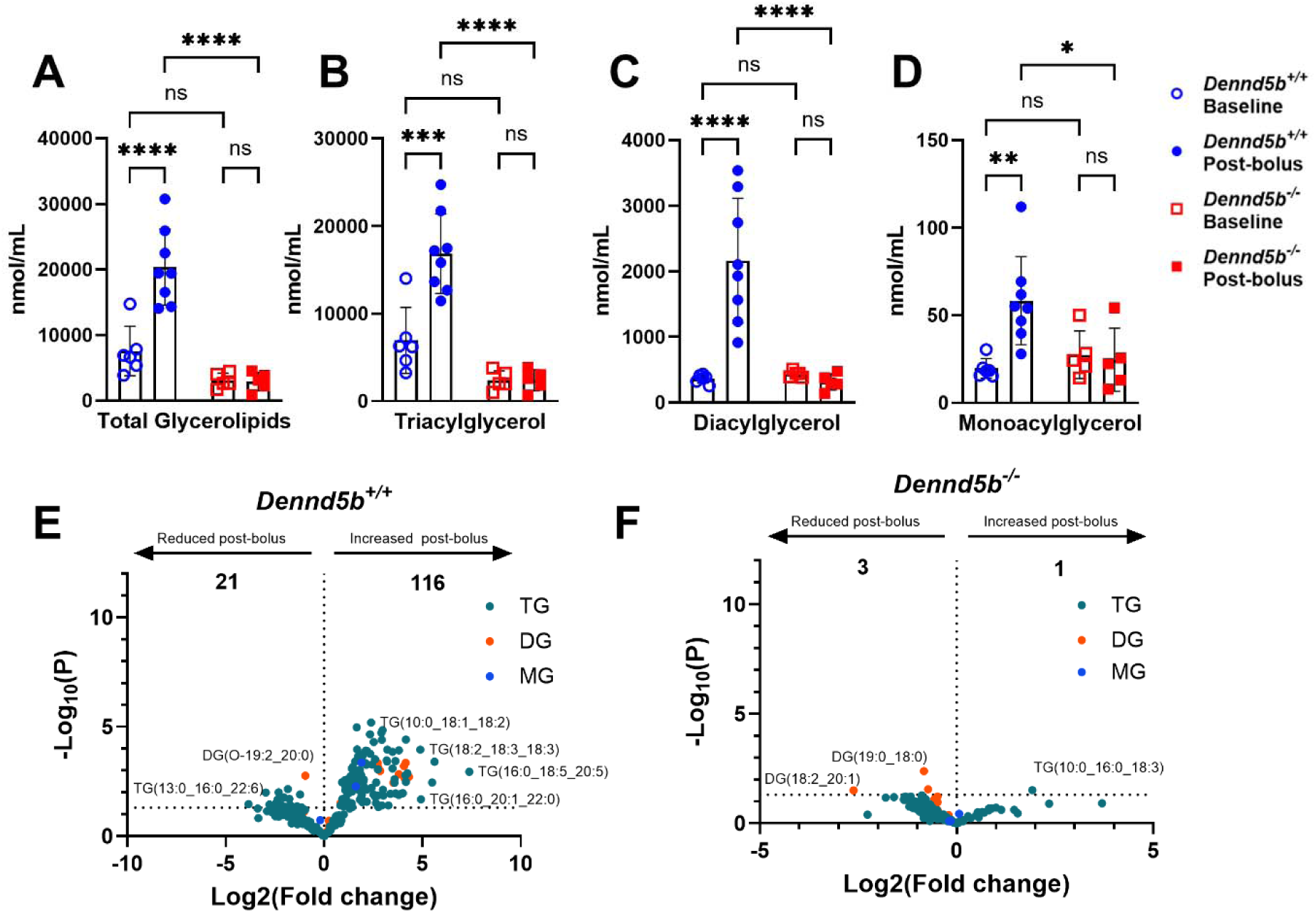
Lymphatic glycerolipids are elevated post-bolus in *Dennd5b^+/+^* but not *Dennd5b^-/-^* lymph. (A-D) Quantification of total glycerolipids and major glycerolipids classes in mesenteric lymph collected from *Dennd5b^+/+^* and *Dennd5b^-/-^* mice at baseline and post-bolus conditions. Data are presented as mean ± SEM. Comparisons were performed using two-way ANOVA with Tukey’s multiple-comparisons test. (E-F) Volcano plot analysis of individual glycerolipids species in post-bolus mesenteric lymph relative to baseline. Each point represents a single lipid species colored by lipid class: TG (teal), DG (orange), and MG (blue). For all statistical analyses: \**p* < 0.05, \*\**p* < 0.01, \*\*\**p* < 0.001, \*\*\*\**p* < 0.0001, ns = not significant.

#### Fatty acyls

At baseline, total fatty acyls (FA) concentrations in mesenteric lymph were comparable between genotypes **(Figure 4A)**. Neither genotype showed post-bolus change in total free fatty acid (FFA) or acylcarnitine concentrations **(Figure 4B, C)**. When concentrations of individual FFA species were examined, 8 FFA species were robustly elevated post-bolus in *Dennd5b*⁺*/*⁺ mice. Interestingly, these included oleic acid (FFA (18:1)) and several very-long-chain fatty acids (FFA (34:1), FFA (31:0), FFA (32:0), FFA (33:0), FFA (36:1), FFA (36:0), and FFA (35:0)) that were either saturated or monounsaturated **(Figure 4D)**. In contrast, *Dennd5b*⁻*/*⁻ mice exhibited minimal FFA changes, with only FFA (34:1) higher post-bolus and 3 FFAs lower post-bolus **(Figure 4E)**. Volcano plots of acylcarnitine species revealed 10 medium- and long-chain acylcarnitines including C14:0, C16:0, and C14-OH were more abundant at baseline **(Figure 4E)**.

**Figure 4.**
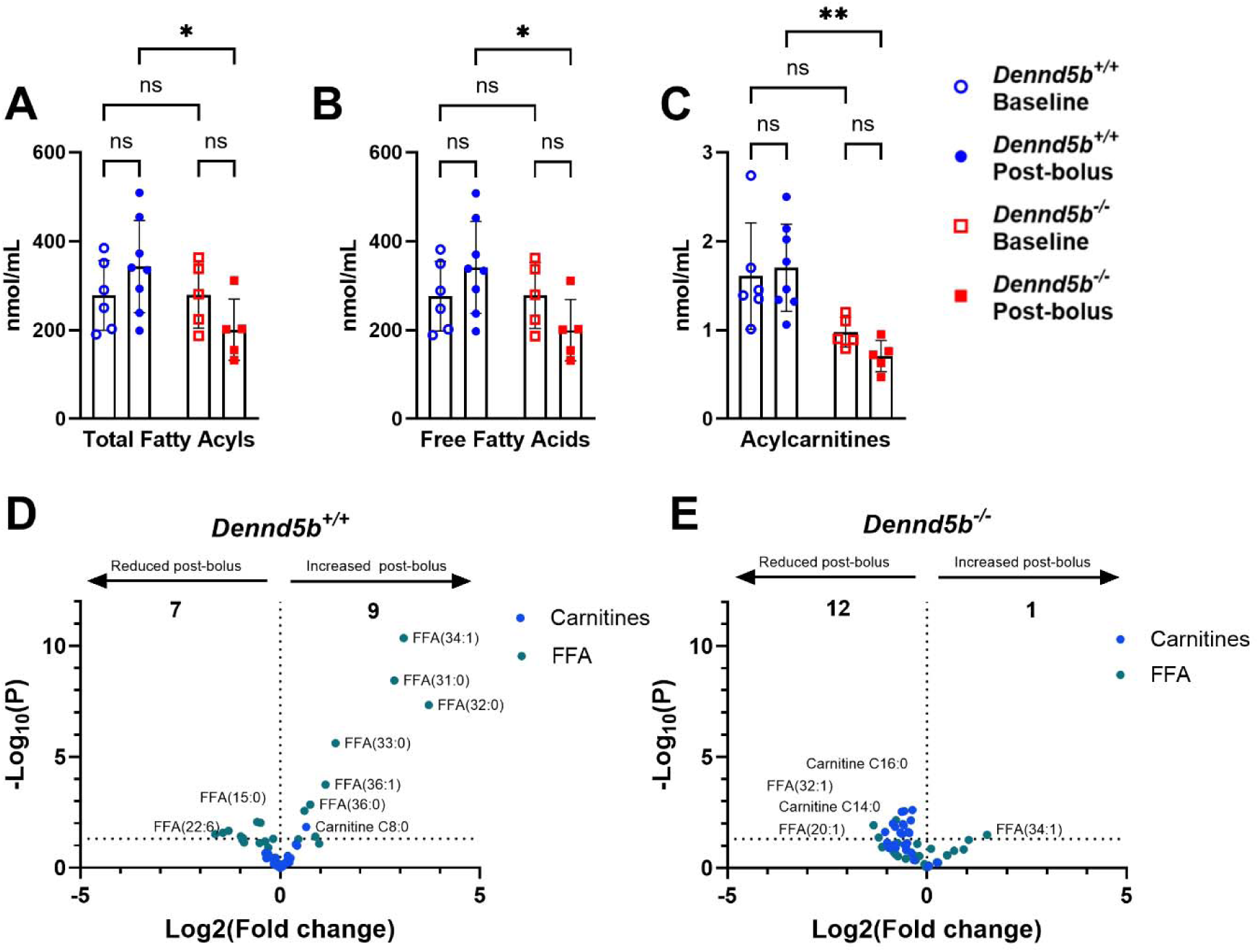
Duodenal lipid bolus increases specific very long-chain free fatty acids in *Dennd5b^+/+^* intestinal lymph. (A-C) Quantification of fatty acyl-containing lipid metabolites in mesenteric lymph collected from *Dennd5b^+/+^* and *Dennd5b^-/-^* mice at baseline and post-bolus conditions. (A) Total fatty acyl concentrations. (B) Free fatty acid (FFA) concentrations. (C) Acylcarnitine concentrations. Comparisons were performed using two-way ANOVA with Tukey’s multiple-comparisons test. (D-E) Volcano plot analysis of individual fatty acyl metabolites in mesenteric lymph following lipid challenge relative to baseline conditions. Each point represents a single lipid species colored by lipid class: free fatty acids (teal) and acylcarnitines (blue). For all statistical analyses: \**p* < 0.05, \*\**p* < 0.01, ns = not significant.

#### Glycerophospholipids

Lymphatic glycerophospholipid composition analysis from *Dennd5b^+/+^* and *Dennd5b^-/-^* lymph samples at baseline revealed that total glycerophospholipid concentration was relatively higher in the *Dennd5b^+/+^* lymph samples compared to *Dennd5b^-/-^* samples but this was not statistically significant **(Figure 5A)**. This observation was similar across glycerophospholipid subclasses **(Figure 5B–D)**. Lipid bolus feeding did not significantly alter total phospholipid content of the mesenteric lymph in either genotype, although in the post-bolus samples total glycerophospholipid concentration was significantly higher in *Dennd5b^+/+^*lymph compared to *Dennd5b^-/-^* **(Figure 5A)**. Analysis of individual glycerophospholipid classes revealed that *Dennd5b^+/+^*mice responded to lipid bolus infusion with increased concentrations of lymphatic phosphatidic acid (PA), phosphatidylethanolamine (PE), and phosphatidylserine (PS) but not phosphatidylcholine (PC), phosphatidylglycerol (PG), and phosphatidylinositol (PI). However, lymph samples from *Dennd5b^-/-^*mice revealed consistently lower concentrations at baseline and a blunted response to the lipid bolus **(Figure 5B–D)**. Interestingly, Lysophosphatidylcholine (LPC) and lysophosphatidylethanolamine (LPE) were readily detectable but not different between *Dennd5b^+/+^* and *Dennd5b^-/-^* mice at both baseline and post lipid feeding conditions **(Supplementary Figure 1)**. In contrast, lysophosphatidylglycerol (LPG) and lysophosphatidylinositol (LPI) were present at very low levels across both genotypes and feeding conditions **(Supplementary Figure 1)**. To examine changes in individual phospholipid species in response to lipid bolus in both genotypes, a fold-change analysis was performed for post-bolus vs baseline concentrations in *Dennd5b^+/+^* and *Dennd5b^-/-^*lymph samples **(Figure 5E–F)**. This analysis revealed that *Dennd5b^+/+^*mice responded to the lipid bolus with increased concentrations of 62 phospholipid species across multiple phospholipid subclasses including PC, PE, and PG **(Figure 5E)**. In contrast, *Dennd5b^- /-^* mice responded to lipid feeding with an increase in concentration of only one phospholipid species PE (20:1_20:5) in the mesenteric lymph **(Figure 5F)**. Interestingly, *Dennd5b^-/-^* mice had several phospholipid species that were lower at post-bolus (n = 27) including LPC and LPE species **(Figure 5F)**. Overall, these results indicate that *Dennd5b* disruption results in significantly impaired lipid feeding induced phospholipid concentration changes in the mesenteric lymph. This is true amongst broader phospholipid subclasses as well as individual phospholipid species. These observations also suggest a potential role of *Dennd5b* in influencing baseline lymphatic phospholipid abundance.

**Figure 5.**
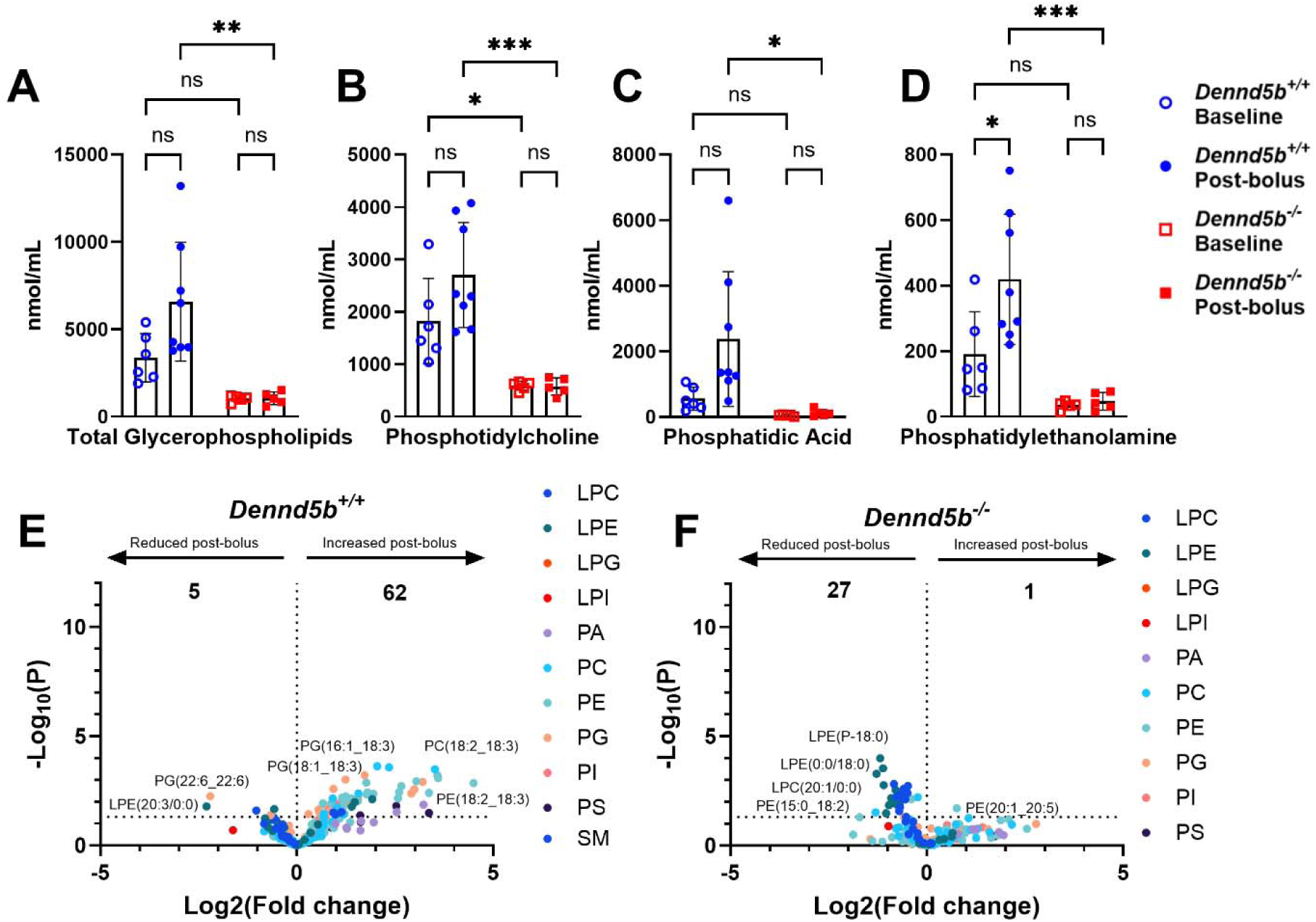
Lymphatic glycerophospholipid species are elevated post-bolus in *Dennd5b^+/+^*but not *Dennd5b^-/-^* lymph. (A-D) Glycerophospholipid analysis of mesenteric lymph samples collected from *Dennd5b^+/+^* and *Dennd5b^-/-^* mice at baseline and post-bolus. (A) Total glycerophospholipid concentrations. (B) Phosphatidylcholine (PC) concentrations. (C) Phosphatidic acid (PA) concentrations. (D) Phosphatidylethanolamine (PE) concentrations. Data are presented as mean ± SEM. Comparisons were performed using two-way ANOVA with Tukey’s multiple-comparisons test. (E-F) Volcano plot analysis of individual glycerophospholipid species in mesenteric lymph following lipid bolus infusion compared to baseline. Each point represents a single lipid species and is colored according to lipid class. For all statistical analyses: \**p* < 0.05, \*\**p* < 0.01, \*\*\**p* < 0.001, ns = not significant.

#### Sphingolipids

At baseline, total sphingolipid concentration was relatively higher in *Dennd5b*⁺*/*⁺ compared to *Dennd5b^-/-^* lymph samples **(Figure 6A)**. Administration of the lipid bolus increased total sphingolipid abundance in the mesenteric lymph of *Dennd5b^+/+^* mice whereas total sphingolipids levels remained unchanged in post-bolus *Dennd5b^-/-^*samples **(Figure 6A)**. Analysis of ceramide subclasses revealed that ceramide species containing saturated fatty acyl chains (Cer-AS), non-hydroxy fatty acids (Cer-NS), alpha-hydroxy fatty acids (Cer-AP), and non-hydroxy fatty acid phytosphingosine (Cer-NP) were the primary contributors to the post-bolus increase in wild type lymph **(Figure 6B-E)**. Cer-AS, and Cer-NP levels were significantly increased (p<0.0001) following lipid bolus infusion in *Dennd5b^+/+^* lymph but Cer- NS and Cer-AP levels did not change in response to lipid infusion **(Figure 6C, D)**. In contrast, *Dennd5b^-/-^* lymph samples exhibited consistently low levels across all ceramide subclasses, with minimal or no response to lipid bolus infusion **(Figure 6B-F)**. Among hexosylceramides (HexCer), only HexCer-NS was significantly elevated in the post-bolus lymph of *Dennd5b^+/+^*mice compared to baseline lymph **(Figure 6G, H)**. In contrast, both HexCer-NS and HexCer- AP species were present at relatively low levels and did not exhibit post-bolus changes in *Dennd5b^-/-^* lymph **(Figure 6G, H)**. To examine changes in individual ceramide species, a fold change analysis was performed comparing post-bolus to baseline concentrations in the lymph **(Figure 6I, J)**. In *Dennd5b^+/+^* mice, lipid administration resulted in an increase in the concentration of a large number of ceramide and hexosylceramides species (n = 91). These increases were observed across several ceramide subclasses, indicating a broad response in sphingolipid concentrations after administration of the lipid bolus. In contrast, *Dennd5b^-/-^* samples showed a severely blunted response, with only one species increased post-bolus **(Figure 6I, J)**. Total sphingomyelin concentrations in lymph did not change in response to the lipid bolus for either genotype **(Supplementary Figure 2A).**

**Figure 6.**
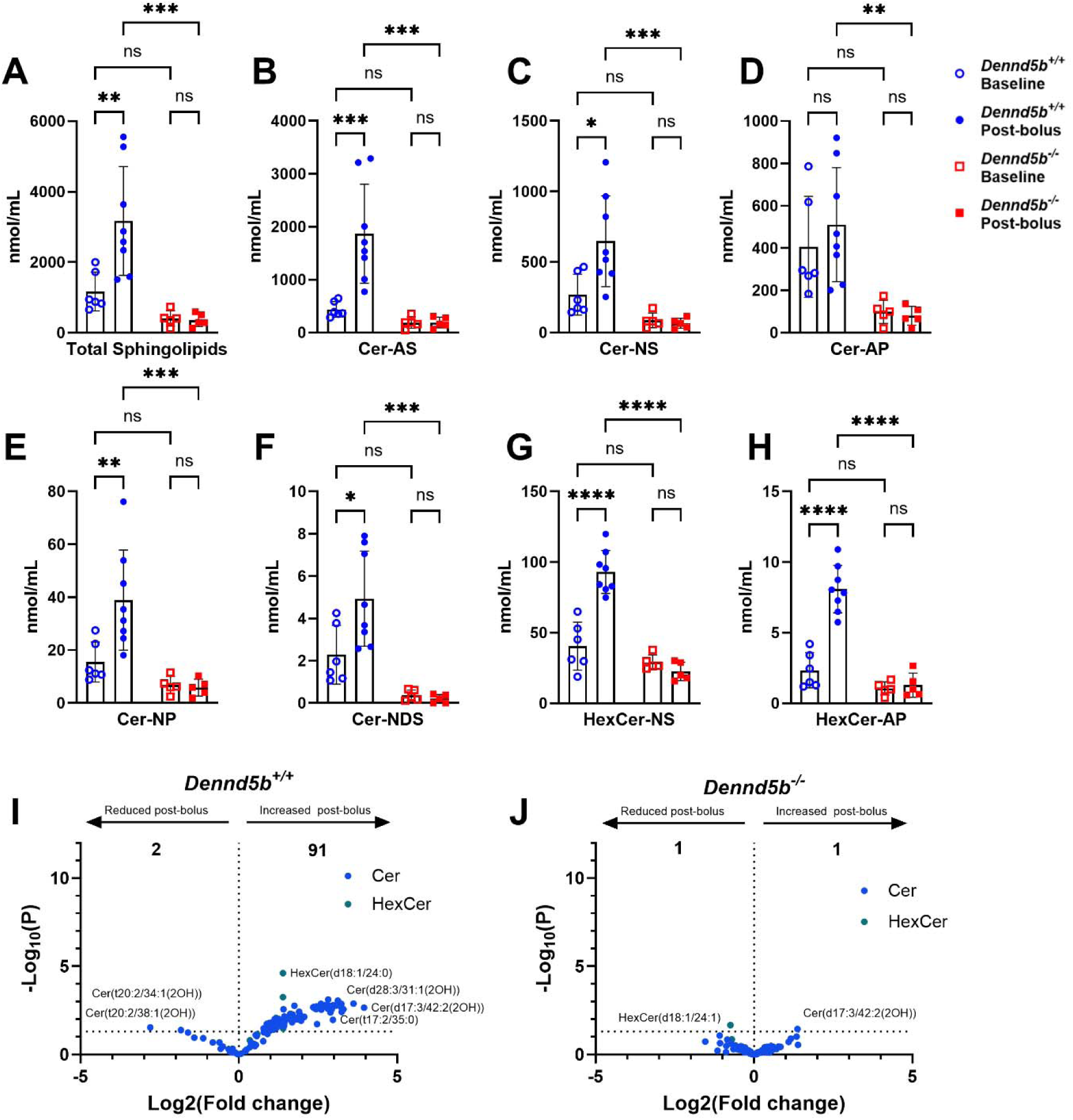
*Dennd5b* deficiency impairs sphingolipid secretion into the mesenteric lymph following duodenal lipid bolus. (A-H) Lipidomic analysis of sphingolipids in mesenteric lymph collected from *Dennd5b^+/+^* and *Dennd5b^-/-^* mice at baseline and post-bolus. (A) Total sphingolipids. (B-F) Ceramide subclasses including Cer-AS, Cer-NS, Cer-AP, Cer-NP, and Cer- NDS. (G-H) Hexosylceramide NS (HexCer-NS), and Hexosylceramide AP (HexCer-AP). Data are mean ± SEM. Comparisons were performed using two-way ANOVA with Tukey’s multiple-comparisons test. (I-J) Volcano plot analysis of sphingolipid species showing individual ceramide (Cer) or hexosylceramide (HexCer) species. For all statistical analyses: \**p* < 0.05, \*\**p* < 0.01, \*\*\**p* < 0.001, \*\*\*\**p* < 0.0001, ns = not significant.

#### Sterol Lipids & Prenol Lipids

Total sterol lipid concentration in mesenteric lymph was higher at baseline in *Dennd5b^+/+^* mice compared to *Dennd5b^-/-^* mice **(Figure 7A)**. In contrast to other lipid classes, total lymphatic sterol lipid concentrations trended downward in response to the lipid bolus in both genotypes **(Figure 7A)**. When sterol lipid subclasses were examined, a similar pattern was observed in both *Dennd5b^+/+^* and *Dennd5b^-/-^* mice for free cholesterol levels **(Figure 7B)**. Cholesteryl esters were significantly higher at baseline in *Dennd5b^+/+^* mice compared to *Dennd5b^-/-^*mice (p < 0.05) **(Figure 7C)**. These levels were dramatically reduced post-bolus in *Dennd5b^+/+^* lymph and unaffected in *Dennd5b^-/-^* lymph **(Figure 7C)**. When analyzed for total bile acid concentration in the mesenteric lymph, *Dennd5b^+/+^* mice had an elevated level at baseline compared to *Dennd5b^-/-^* mice **(Figure 7D)**. Both *Dennd5b^+/+^* and *Dennd5b^-/-^* mice showed minimal response to changes in total bile acid concentration in the mesenteric lymph in response to the lipid bolus **(Figure 7D)**. The observed differences in total bile acid concentrations among *Dennd5b^+/+^*and *Dennd5b^-/-^* mice were mostly contributed by the differences in concentrations of taurochenodeoxycholic acid **(Figure 7E)**. All other bile acids detected were present at minimal concentrations at baseline and post lipid feeding in both *Dennd5b^+/+^*and *Dennd5b^-/-^* mice **(Figure 7F-H)**. Volcano plot analysis revealed that in *Dennd5b^+/+^* mice, two individual sterol species including CE (18:2) and CE (16:0) were reduced post-bolus but no sterols were found to be enriched after lipid feeding **(Figure 7I)**. In addition, the concentrations two individual bile acid species including glycocholic acid and lithocholic acid-3-sulfate were increased post-bolus in *Dennd5b^-/-^* mice **(Figure 7J)**. Similar to bile acids, lipidomic analysis revealed that Coenzyme Q was preset at very low concentrations in the mesenteric lymph across genotypes and feeding conditions **(Supplementary Figure 2B)**.

**Figure 7.**
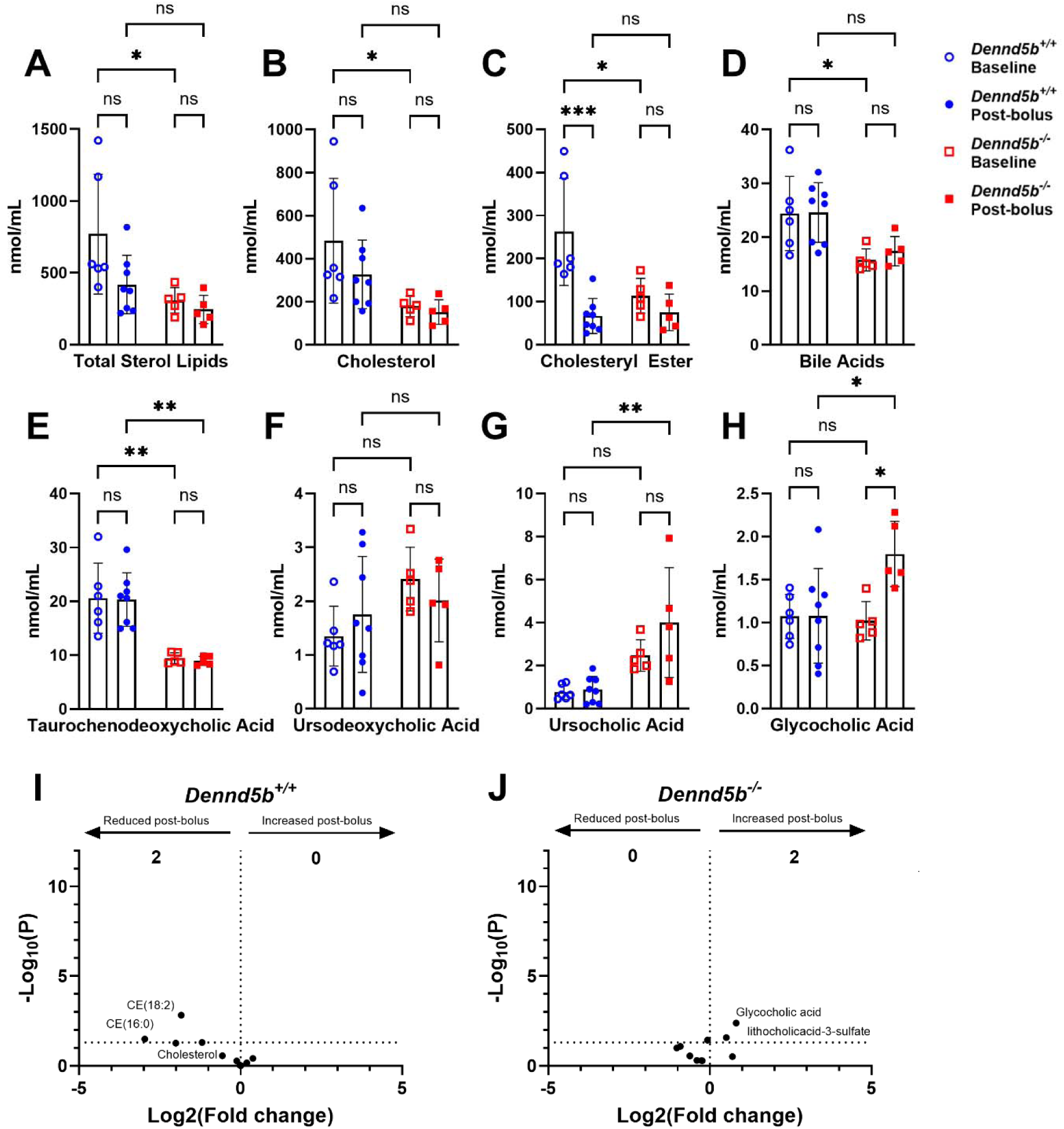
*Dennd5b* deficiency has minimal impact on lymphatic sterol concentrations after duodenal lipid bolus. (A-H) Quantification of sterol lipids in mesenteric lymph collected from *Dennd5b^+/+^* and *Dennd5b^-/-^* mice at baseline and post-bolus conditions. (A) Total sterol lipids. (B) Cholesterol. (C) Cholesteryl esters. (D) Bile acids. (E) Taurochenodeoxycholic acid. (F) Ursodeoxycholic acid. (G) Ursocholic acid. (H) Glycocholic acid. Data are mean ± SEM. Comparisons were performed using two-way ANOVA with Tukey’s multiple-comparisons test. (I-J) Volcano plot analysis of individual sterol species in mesenteric lymph following lipid bolus infusion relative to baseline. For all statistical analyses: \**p* < 0.05, \*\**p* < 0.01, \*\*\**p* < 0.001, ns = not significant.

## Discussion

In this study, we performed a detailed analysis of the lipid composition of mesenteric lymph before and after administration of a duodenal lipid bolus. These samples were obtained using a conscious lymph fistula model that enabled collection of mesenteric lymph, before dietary lipids are exposed to modifying factors in plasma. Our goals were to characterize the changes to the mesenteric lymph lipidome during the postprandial state and to test the hypothesis that *Dennd5b* deficiency impairs the absorption of dietary lipids beyond the previously described impairment of triglyceride absorption (16). Three major findings emerged from this study. First, the categorical lipid composition of post-bolus lymph in wild type mice generally shifted toward that of the administered lipid bolus. Second, sphingolipids were a notable exception, exhibiting a disproportionate increase in post-bolus lymph. Third, *Dennd5b* deficiency broadly suppressed the appearance of dietary lipids in mesenteric lymph, with little or no post-bolus increase across most major lipid categories and species.

Administration of a duodenal lipid bolus to wild type mice resulted in significant intestinal output of total lipid to the mesenteric lymph and a shift in the lymph lipidome composition toward the composition of the administered Intralipid. These findings suggest that, at the level of broad lipid categories, the composition of newly secreted intestinal lipoproteins is strongly influenced by the composition of the ingested lipid and that many ingested lipids maintain their categorical identity during absorption. While this is generally expected, this relationship is notable because many dietary lipids undergo extensive hydrolysis before uptake by enterocytes, intracellular reesterification, lipoprotein packaging, and secretion. Thus, there is potential for reesterification of lipids into forms that are distinct from the ingested material.

For six of the seven lipid categories analyzed, the direction of the post-bolus change in wild type lymph was consistent with the relative abundance of that lipid category in the administered lipid compared with baseline lymph (Figure 2). Sterols (ST), glycerophospholipids (GP), fatty acyls (FA), prenol lipids (PR), and other lipids were less abundant in the administered lipid than in baseline lymph and consequently decreased in relative molar abundance following the bolus. Conversely, glycerolipids (GL), which were more abundant in Intralipid than in baseline lymph, increased after bolus administration. Sphingolipids (SP) were the notable exception. Although their relative abundance was lower in Intralipid than in baseline lymph, their molar percentage increased following bolus administration. This disproportionate increase suggests that intestinal processing of a mixed lipid bolus may stimulate sphingolipid synthesis, mobilization, and/or secretion into mesenteric lymph. Alternatively, sphingolipid appearance in lymph may follow absorption and secretion kinetics that differ from those of other lipid classes. Because only a single post-bolus interval was examined, these possibilities cannot be excluded. Overall, the lipid composition of postprandial mesenteric lymph in wild type mice largely reflected the composition of the administered lipid, with sphingolipids displaying a distinct response at the time point analyzed.

In striking contrast to wild type mice, *Dennd5b^-/-^* mice exhibited essentially no shift in the relative molar composition of any lipid category in response to the lipid bolus. Lipidomic analysis demonstrated that loss of *Dennd5b* impairs absorption not only of TG, as previously reported, but also of most of the lipid species analyzed. In the post-bolus state, wild type mice experience the most significant increases (by concentration fold-change) in glycerolipids (TG, DG, and MG), glycerophospholipids (mostly LNAPE, PC, PE, PS, and PA), and sphingolipids (Cer and HexCer). However, in *Dennd5b^-/-^* lymph, concentrations of all of these lipids either did not respond or had a markedly reduced response to the lipid bolus. Together, these findings support the conclusion that *Dennd5b* is required for the normal postprandial export of a broad range of lipid species into mesenteric lymph and support its essential role in the secretion of lipid-loaded intestinal chylomicrons.

Differences between genotypes were evident even under baseline conditions. Wild-type and *Dennd5b^-/-^*mice displayed distinct baseline lymph lipidomes, with lower concentrations of several lipid species in *Dennd5b^-/-^* mice, including TG, PC, cholesterol, cholesterol ester, and taurochenodeoxycholic acid. These differences suggest that the influence of *Dennd5b* on intestinal lipid homeostasis is not restricted to the acute response to a lipid bolus. Rather, *Dennd5b* deficiency may alter basal lipid trafficking, secretion, or enterohepatic lipid metabolism even in the absence of an acute dietary lipid challenge.

The profoundly blunted postprandial secretion of glycerolipids in *Dennd5b^-/-^* mice is consistent with our previous reports demonstrating impaired secretion of lipid-loaded chylomicrons (16, 18). The lower abundance of phospholipids and sphingolipids in post-bolus *Dennd5b^-/-^* lymph further suggests that delivery of these lipids to mesenteric lymph is tightly coupled to intestinal chylomicron secretion. The sphingolipid phenotype is particularly noteworthy because wild-type mice exhibited a disproportionate increase in sphingolipids following the lipid bolus. This increase may reflect active intestinal sphingolipid synthesis as a contributing source alongside direct absorption of sphingolipids present in the lipid bolus. Enterocytes can generate ceramides through *de novo* synthesis from serine and fatty acyl-CoAs, particularly palmitoyl-CoA, and increased dietary fatty acid availability can enhance flux through this pathway (20, 21). Consistent with this mechanism, high-fat feeding has been shown to increase intestinal ceramide production, including C16:0 ceramide, and promote incorporation of intestinally derived ceramides into chylomicrons for transport through the mesenteric lymph (21). Ceramide also serves as a central precursor for more complex sphingolipids, including sphingomyelins and glycosphingolipids such as HexCer, providing a potential mechanism by which an acute lipid load could alter multiple sphingolipid species within enterocytes before secretion (20, 22). Dietary sphingolipids themselves are extensively hydrolyzed in the intestinal lumen, with only a fraction of absorbed sphingoid bases reincorporated into enterocyte ceramides and complex sphingolipids (23, 24). Thus, the increase in ceramides and HexCer observed in post-bolus lymph may represent a combination of intestinal synthesis and remodeling stimulated by dietary fatty acid availability, followed by incorporation of these newly generated sphingolipids into nascent chylomicrons. The near absence of this response in *Dennd5b^-/-^* mice is therefore consistent with impaired chylomicron secretion preventing the normal postprandial export of enterocyte-derived sphingolipids into the mesenteric lymph. Alternatively, *Dennd5b* deficiency could indirectly alter enterocyte sphingolipid synthesis, remodeling, or trafficking as a consequence of intracellular lipid retention. Additional studies would be required to determine whether the sphingolipid phenotype is solely secondary to impaired chylomicron secretion or reflects an additional role for *Dennd5b* in intestinal sphingolipid metabolism.

Total free fatty acids, as a lipid category, did not significantly increase in post-bolus lymph of wild type mice. Analysis of individual fatty acid species, however, revealed significant elevation of several specific free fatty acids in post-bolus lymph. These consisted predominantly of saturated and monounsaturated very-long-chain fatty acids with acyl chains exceeding 30 carbons. Many of these species were also detected as free fatty acids in the administered Intralipid, consistent with a dietary origin. Their appearance in mesenteric lymph raises questions regarding how these unusually long-chain species are absorbed and transported by the intestinal epithelium. It remains unclear whether they traverse enterocytes as free fatty acids or undergo transient esterification and subsequent hydrolysis during absorption and secretion.

An important strength of this study was the use of the conscious lymph fistula model, which enabled collection of mouse mesenteric lymph and characterization of newly secreted intestinal lipids before chylomicrons undergo extensive remodeling in the systemic circulation. Combining this approach with comprehensive lipidomic profiling allowed us to detect lipid changes that would likely have been missed by measurements performed on isolated plasma chylomicrons. Several limitations should be considered. First, our lipidomic analysis was limited to baseline and one postprandial time point (between 2-3 hours post-bolus) and therefore does not capture the full picture of lipid absorption kinetics. Second, lipidomics analysis was performed on whole lymph, not isolated chylomicrons, therefore, the current analysis does not establish the physical association of individual lipid species specifically with chylomicrons. Although most of the lipids analyzed were expected to be associated with chylomicrons, especially in the post-bolus state, a relatively minor amount of lipid from other lipoproteins or free lipids may be present in the mesenteric lymph. Another limitation is that this study only examined lymph from male mice. Given the known sex-differences in postprandial lipid absorption kinetics in humans and mice (18, 25, 26), it is plausible that biological sex may impact postprandial lymph lipidome. Last, the intralipid bolus, lacking substantial protein and carbohydrate, does not recapitulate the nutrient composition of a common meal. Future studies using mixed-nutrient meals and multiple postprandial time points will be important for determining how dietary composition influences the kinetics and composition of lipid transport into mesenteric lymph.

### Conclusions

In this study, we provide a detailed lipidomic characterization of mesenteric lymph following intestinal lipid administration in wild type and *Dennd5b^-/-^* mice. In wild type mice, the postprandial lymph lipidome generally shifted toward the composition of the administered lipid, indicating a strong relationship between dietary lipid composition and the lipids exported to the intestinal lymphatics. Sphingolipids were a notable exception, exhibiting a greater than expected postprandial increase that suggests distinct regulation of their intestinal synthesis, processing, or secretion.

Our findings also substantially expand the lipid phenotype associated with *Dennd5b* deficiency. The present study demonstrates that the effect of genetic *Dennd5b* disruption extends well beyond triglyceride, resulting in broadly impaired postprandial appearance of multiple lipid classes in mesenteric lymph. These results support a model in which *Dennd5b*-dependent chylomicron secretion serves as a major pathway for the coordinated intestinal export of a diverse set of dietary and enterocyte-derived lipids.

Because chylomicrons transport a complex mixture of bioactive lipids in addition to triglycerides, alterations in their abundance and composition may have consequences beyond energy delivery. The lipid composition of newly secreted chylomicrons could influence particle metabolism, tissue lipid delivery, cellular signaling, and ultimately cardiometabolic disease. Determining whether specific changes in the lipid composition of intestine-derived lipoproteins contribute to systemic metabolic health and atherosclerotic cardiovascular disease represents an important direction for future investigation.

## Data availability statement

The data that support the findings of this study are available on request from the corresponding author.

## Acknowledgments and grant support

This work was supported by the National Institutes of Health National Institute of Digestive, Diabetes and Kidney Diseases, R01DK133184 (S.M.G.). The content is solely the responsibility of the authors and does not necessarily represent the official views of the National Institutes of Health.

