## Supplemental Figures for "Lipidomic analysis of postprandial mesenteric lymph from wild type and *Dennd5b* knockout mice"

*** Corresponding Author:**

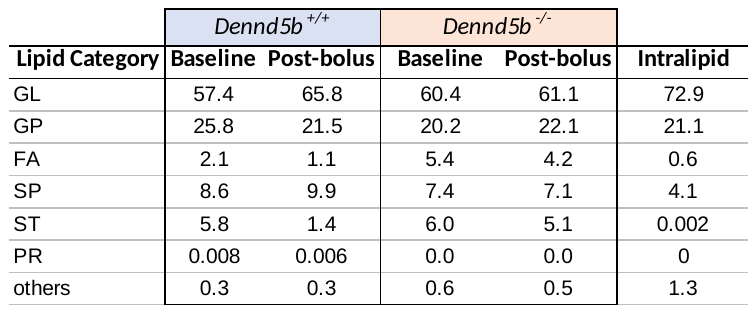

**Supplementary Table 1. Molar percent composition of baseline and post-bolus lymph from wild type and *Dennd5b^-/-^* mice.**

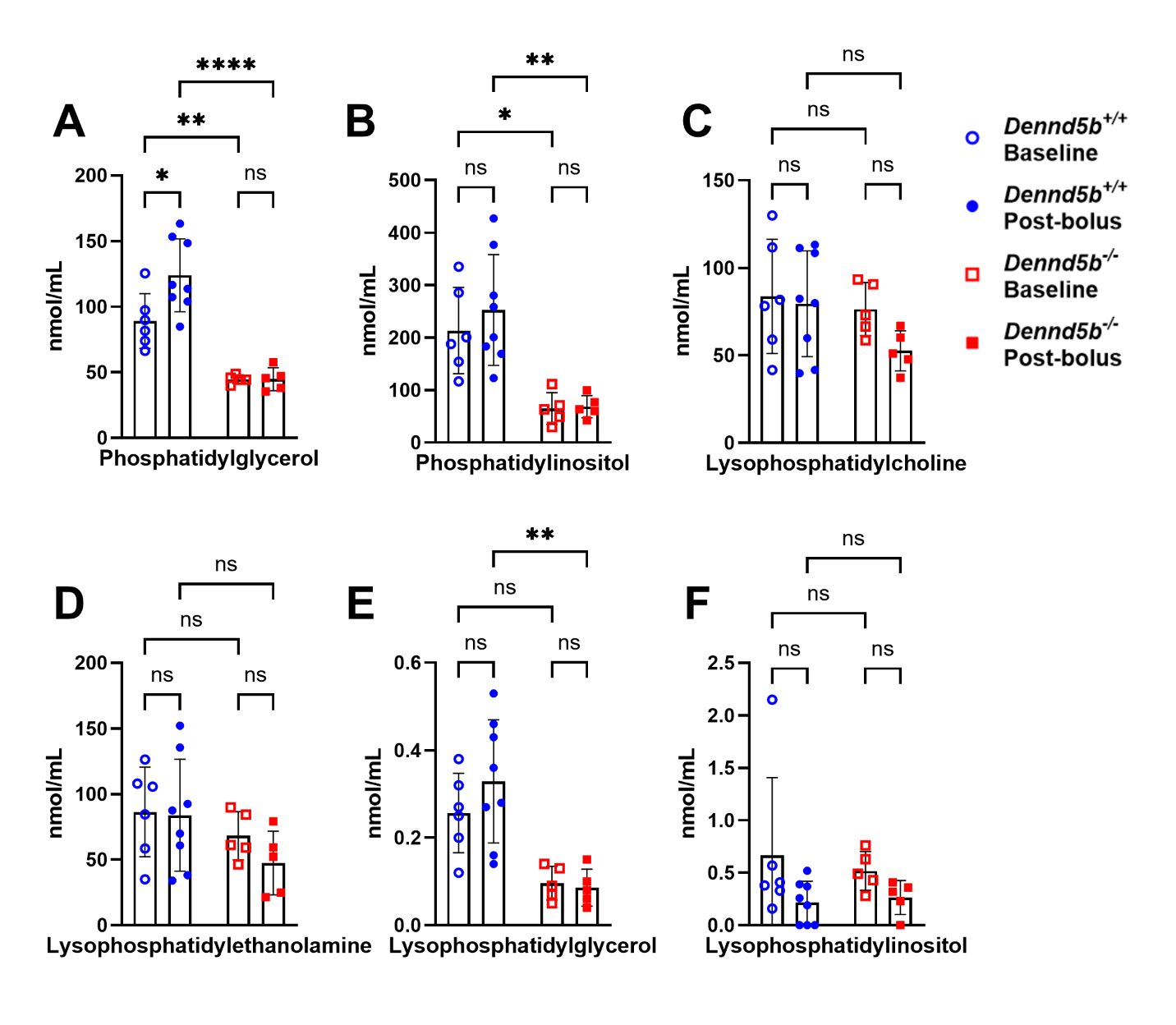
**Supplementary Figure 1. Changes in concentration of glycerophospholipids and lysophospholipids in the mesenteric lymph of Dennd5b^+/+^ and Dennd5b^-/-^ mice at baseline and following lipid bolus infusion.** (A-D) Glycerophospholipids and lysophospholipids analysis of mesenteric lymph samples collected from Dennd5b^+/+^ and Dennd5b^-/-^ mice at baseline and post-bolus. (A) Phosphatidylglycerol (B) Phosphatidylinositol (C) Lysophosphatidylcholine (D) Lysophosphatidylethanolamine (E) Lysophosphatidylglycerol (F) Lysophosphatidylinositol concentrations in mesenteric lymph. Data are presented as mean ± SEM. Comparisons were performed using two-way ANOVA with Tukey's multiple-comparisons test. For all statistical analyses: **p* < 0.05, ***p* < 0.01, *****p* < 0.0001; ns, not significant.

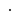

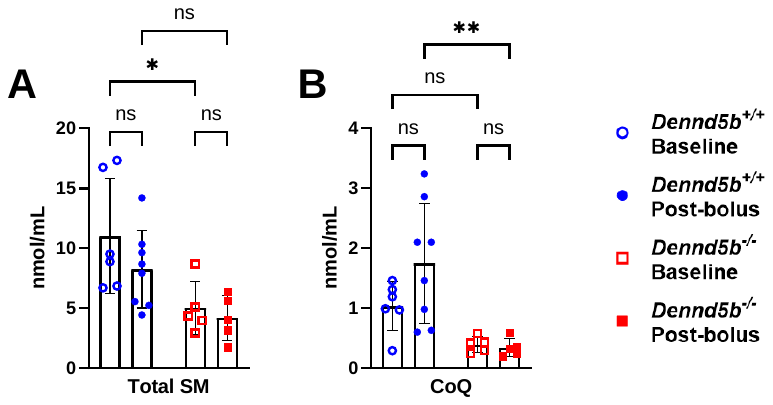

**Supplementary Figure 2. Changes in concentration of total sphingomyelin and coenzyme Q in the mesenteric lymph of Dennd5b^+/+^ and Dennd5b^-/-^ mice at baseline and following lipid bolus infusion**. Quantification of (A) total sphingomyelin (SM) and (B) coenzyme Q (CoQ) in mesenteric lymph. Data are presented as mean ± SEM. Comparisons were performed using two-way ANOVA with Tukey's multiple-comparisons test. For all statistical analyses: **p* < 0.05, ***p* < 0.01; ns, not significant.
